# Loop-closure Kinetics Reveal a Stable, Right-handed DNA Intermediate in Cre Recombination

**DOI:** 10.1101/695130

**Authors:** Massa J. Shoura, Stefan M. Giovan, Alexandre V. Vetcher, Riccardo Ziraldo, Andreas Hanke, Stephen D. Levene

## Abstract

In Cre site-specific recombination, the synaptic intermediate is a recombinase homotetramer containing a pair of DNA target sites. The strand-exchange mechanism proceeds via a Holliday-junction (HJ) intermediate; however, the geometry of the DNA segments in the synapse has remained highly controversial. In particular, all crystallographic structures are consistent with an achiral planar Holliday-junction (HJ) structure, whereas topological assays based on Cre-mediated knotting of plasmid DNAs are consistent with a right-handed chiral junction. Here we use the kinetics of loop closure involving closely spaced (131-151 bp), directly repeated loxP sites to investigate the *in-aqueo* ensemble of conformations for the longest-lived looped DNA intermediate. Fitting the experimental site-spacing dependence of the loop-closure probability, *J*, to a statistical-mechanical theory of DNA looping provides evidence for substantial out-of-plane HJ distortion. This result unequivocally stands in contrast to the square-planar intermediate geometry determined from crystallographic data for the Cre-loxP system and other int-superfamily recombinases. *J* measurements carried out with an isomerization-deficient Cre mutant suggest that the apparent geometry of the wild-type complex may result from the temporal averaging of diverse right-handed and achiral structures. Applied to Cre recombinase, and other biological systems, our approach bridges the static pictures provided by crystal structures and the natural dynamics of macromolecules *in vivo*. This approach thus advances a more comprehensive dynamic analysis of large nucleoprotein structures and their mechanisms.

## Introduction

Changes in DNA topology, manifested by properties such as supercoiling, knotting, and catenation, are integral to numerous cellular processes including DNA replication, transcription, and recombination.^1–7^ Using circular DNA as a model system to investigate these processes allows associated topological changes to be readily characterized by established biophysical techniques such as gel electrophoresis.^8–10^ However, making connections between changes in topological parameters and perturbations of canonical DNA geometry is not straightforward for several reasons. First, geometric solutions consistent with a given topology are rarely unique. This is the case with DNA supercoiling, for which the sum of DNA twist and writhe is conserved (and equal to the linking number), but not the individual values of those parameters. Partly because of this ambiguity, relating enzyme-mediated changes in DNA topology to DNA geometry often requires some assumptions about the geometries of nucleoprotein intermediates in the enzymatic pathway.^11,12^ These assumptions can be difficult to confirm independently through measurements in solution; moreover, atomic-resolution crystallographic structures are not always helpful in resolving structural ambiguities because of inherently limited information about conformational dynamics. Thus, nuances such as conformational changes in the context of a cellular environment may not be fully revealed. Finally, in order to engineer molecular conformations having specific functions, for example construction of specific DNA topologies,^13–15^ there is value in knowing the precise configuration (position in space, orientation, and chirality) of DNA segments bound to the enzyme and also information about the conformational dynamics of the nucleoprotein structure

The Cre-recombination system of bacteriophage P1 has become an important tool for the genetic manipulation of higher organisms^16^ and is a paradigm for site-specific DNA-recombination mechanisms employed by the λ-integrase (λ-int) superfamily of recombinases.^17, 18^ In the life cycle of phage P1 Cre plays a critical role in genome maintenance by unlinking newly replicated sister chromosomes to ensure their proper segregation following DNA replication.^19^ Cre acts at a 34-bp wild-type DNA recombination-target sequence called loxP, which consists of 13-bp perfect inverted repeats flanking a non-palindromic 8-bp core region^20^ and defines the site’s absolute orientation. On a linear DNA molecule Cre-mediated recombination involving a pair of tandemly repeated loxP sequences generates a pair of deletion products (one linear, one circular), each bearing a single copy of loxP (Figure 1). DNA-cleavage steps involve a phosphotyrosine intermediate similar to those employed by type-IB topoisomerases.^21^ As with other members of the λ-int superfamily, recombination takes place via two successive rounds of DNA-strand cleavage and exchange steps that respectively generate and resolve a four-stranded, Holliday-junction (HJ) DNA intermediate.^20, 22^

**Figure 1.**
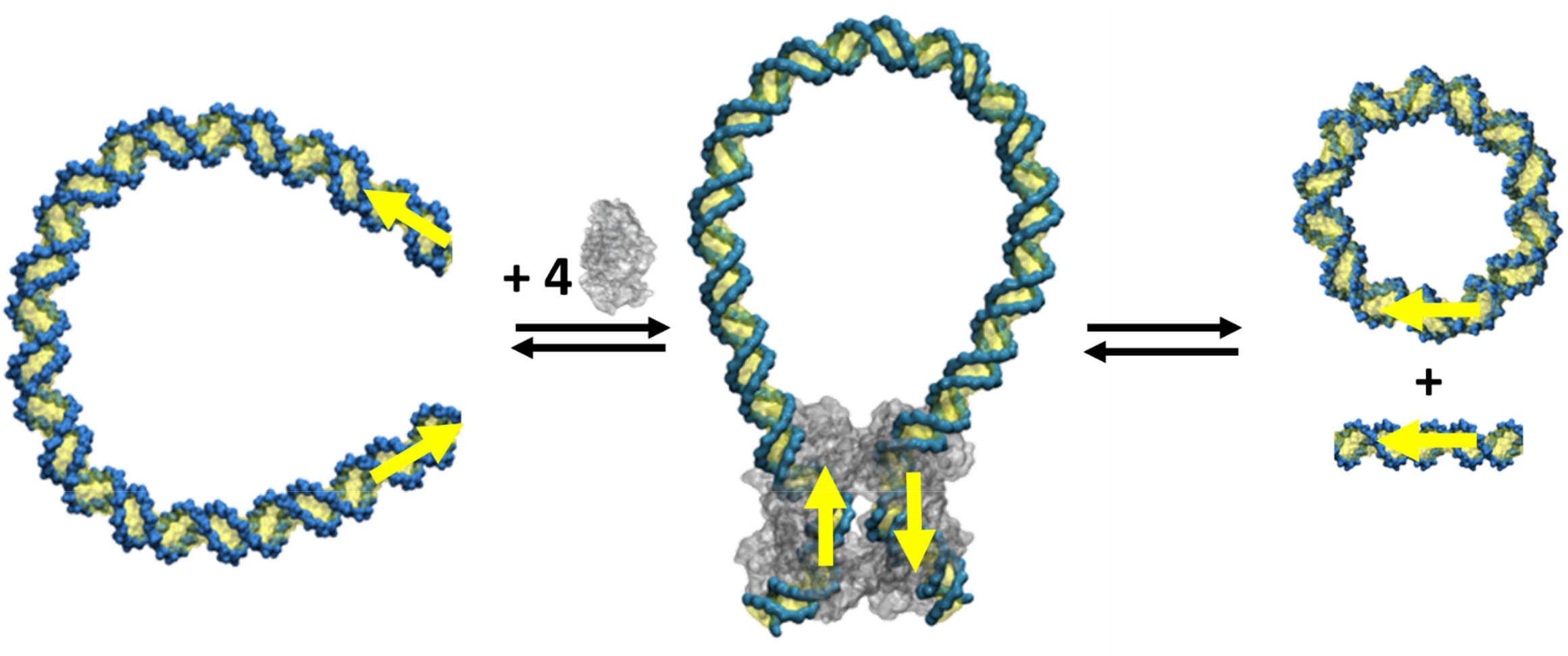
Products generated by Cre recombinase acting on the DNA molecules used in this study. In the first step DNA fragments flanked by directly repeated loxP sites (indicated by yellow arrows) form a DNA loop via site synapsis, which is mediated by a Cre-protein tetramer. Two sequential cleavage/strand-exchange events generate a short linear molecule and a DNA circle as recombinant products in the second step.

High-resolution crystal structures of Cre synaptic complexes formed with duplex and junction DNAs provided the first insights into mechanistic details of recombination in these systems. On the basis of these structures, the chemical steps in Cre recombination have been explained in terms of DNA strand exchanges taking place within a twofold-symmetric, nearly square-planar arrangement of the DNA duplexes.^18, 22–28^ There is, however, a major still-unresolved disagreement between the square-planar exchange mechanism and evidence for a chiral recombination intermediate from topological studies of the Cre reaction on circular DNAs. Analyses of the chirality biases in knotted Cre-recombination products (and also those generated by the mechanistically similar yeast recombinase Flp) strongly suggest that the synaptic intermediate involves a right-handed crossing of target sites.^29^ A chirality bias is also supported by atomic-force microscopy (AFM) images of synaptic complexes formed on circular DNAs,^30^ although the handedness of crossings in the complex could not be determined in those experiments. Studies employing tethered single-molecule techniques have sought to address the dynamics of the Cre-loxP synapse in solution,^31–34^ but are potentially subject to various artifacts that arise from the tethering constraint.^35^ Thus, information about the structure of the Cre synaptic complex free in solution, as opposed to the crystal form or another immobilized state, has not been available.

To address this disagreement, we investigated the *in-aqueo* ensemble of conformations for the longest-lived looped DNA intermediate in Cre recombination. We measured rates of intramolecular Cre-loxP recombination for twenty DNA constructs bearing closely spaced (131-151 bp), directly repeated loxP sites. Detailed analysis of the measured J factors for this series of constructs using a statistical-mechanical theory for loop closure yielded a well-defined set of geometric parameters corresponding to the minimum mechanical-energy structure of the Cre-loxP synaptic complex. This structure places the loxP half sites in the recombinase-DNA complex at the base of a looped DNA conformation that has close to the 90-degree internal angle expected for a protein-free HJ. However, the average dihedral angle relating bent DNA segments at the loop ends amounts to a right-handed crossing of approximately 66 degrees, suggesting a substantial departure from planar-intermediate geometry.

Our experiments in solution thereby provide a structural model consistent with the right-handed chirality of Int-superfamily recombination sites inferred from the combination of topological methods and tangle solutions from knot theory.^29^ Applied to Cre recombinase and other biological systems, our approach bridges the static pictures provided by crystal structures and the natural dynamics of macromolecules *in vivo*. This approach thus advances a more comprehensive dynamic analysis of large, biologically active nucleoprotein structures and their mechanisms.

## Results

### Determining Loop-closure Probabilities from Kinetic Data

Independent measurements of inter- and intramolecular recombination rates permit quantitative determination of the probability of loop formation, *J*. Formally, *J* is a quotient of rate constants for intra- and intermolecular synapsis, given by

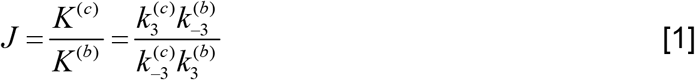

where each of the *k*_3_/*k*_−3_ ratios pertains to corresponding apparent forward and reverse synapsis steps for the loop-closure and bimolecular synapsis reactions (superscripts ^*(c)*^ and ^(b)^, respectively).^36^ Numerical values of the rate constants are fitted parameters in a system of ordinary differential equations evaluated from the time-dependent quenching of a FRET donor signal during the Cre reaction.^36^

An additional challenge of J-factor measurements comes from the observation that inter- and intramolecular reactions generally do not take place under identical conditions. The limited range of intermolecular-recombination conditions is similar to that encountered in ligase-catalyzed bimolecular-joining reactions,^22^ which require higher concentrations of DNA substrate than for the corresponding intramolecular reaction. Failing to take possible dependencies on substrate or enzyme concentration into account can lead to large errors in *K*^(*b*)^, and hence in the absolute value of *J*. The most rigorous approach for obtaining absolute measurements of *J* is to determine individual pairs of apparent rate constants for the intra- and intermolecular reactions under identical solution conditions, extrapolating to intramolecular reaction conditions where necessary. J factors for Cre-mediated looping of loxP sites separated by more than 800 bp have been determined using this method.^36^

In Table 1 we report the *J* values for DNA loops of size *n* = 130 bp to 153 bp (taken as curvilinear distances in base pairs between the centers of directly repeated loxP sites) using the wild-type enzyme. Recombination of the directly repeated loxPs is a deletion reaction that generates a small, circular recombination product whose size in base pairs is exactly equal to that of the Cre-mediated loop along with a 34-bp linear product (Figure 1). The recombination substrates were designed with internal donor- and acceptor-fluorophore labels located within the spacer regions of loxP sites, which are positioned near the ends of the molecules (see Supplementary Figure 1). In order to prevent the formation of intermolecular-recombination products, kinetic assays were carried out at bulk loxP concentrations not exceeding 0.5 nM, an order of magnitude below the target-site bulk-concentration threshold for intermolecular Cre recombination.^36^ Post-hoc analysis of the reactions by agarose-gel electrophoresis also confirmed that the yield of intermolecular-recombination products was negligible under our conditions (data not shown).

**Table 1.** J-factor values and uncertainties expressed as standard deviations, *J_err_*. Each data point is the average of at least five independent measurements.

| Loop size ($n$), bp | $J_{exp}$, $10^{-9}$ moles L$^{-1}$ | $J_{err}$, $10^{-10}$ moles L$^{-1}$ |
| --- | --- | --- |
| 131 | 1.1 | 5.0 |
| 133 | 2.5 | 2.4 |
| 134 | 6.4 | 2.4 |
| 135 | 7.9 | 2.7 |
| 137 | 17.1 | 3.3 |
| 138 | 14.8 | 4.0 |
| 139 | 10.6 | 3.1 |
| 140 | 2.9 | 3.6 |
| 142 | 2.2 | 2.9 |
| 143 | 2.7 | 2.9 |
| 144 | 4.8 | 3.6 |
| 145 | 14.0 | 2.9 |
| 146 | 18.5 | 2.4 |
| 147 | 19.4 | 2.4 |
| 148 | 18.7 | 3.2 |
| 149 | 13.5 | 2.9 |
| 151 | 7.2 | 4.1 |

Figure 2 shows that excellent fits are obtained for the time-dependent FRET signals to ordinary-differential equation (ODE) solutions for the intramolecular single-intermediate Cre-recombination pathway.^36^ A fixed parameter is required in the fitting routine corresponding to the steady-state quenching of donor fluorescence upon synapsis of the donor- and acceptor-labeled loxP sites. This quenching is quantified by the ratio of donor quantum yields in the Cre synaptosome to that for the Cre-bound donor-labeled substrate alone, 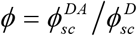, and was taken to be 0.12, a value determined previously for intermolecular synapsis.^36^ The rate constants 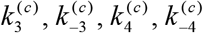, which appear in the system of ODEs modeling the recombination mechanism, were treated as adjustable parameters. Remaining rate constants were fixed at the same values used in previous studies.^36, 37^ As shown in Figure 2, the overall amplitude of the FRET signal varied significantly with loop size, asymptotically approaching decreases of between 10 and 25 percent over the course of 45-minute reactions.

**Figure 2.**
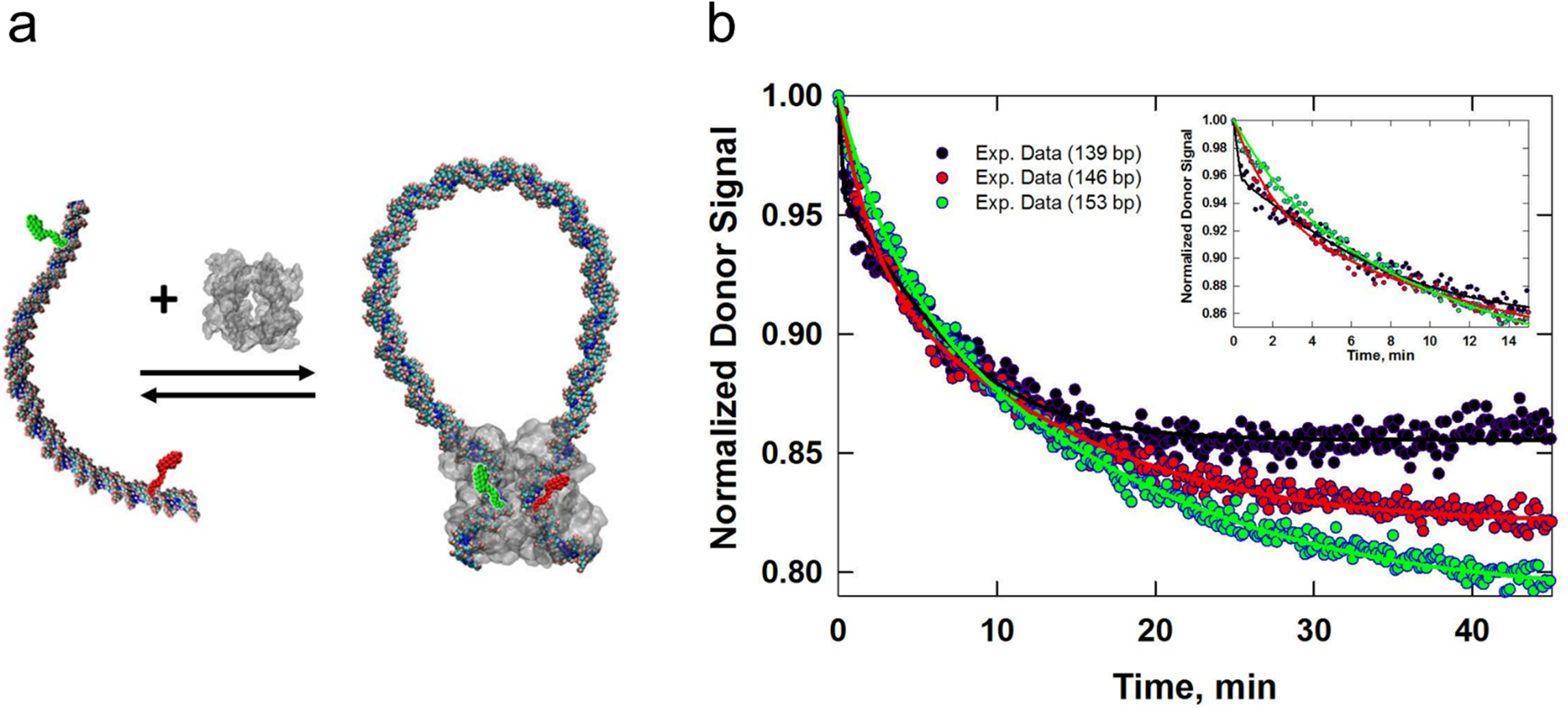
Intramolecular synapsis and recombination kinetics obtained from time-dependent FRET measurements. (**a**) Schematic of the intramolecular reaction carried out on a DNA fragment bearing donor- and acceptor-labeled loxP sites. (**b**) Donor-fluorescence signal, which monitors donor quenching via FRET during site synapsis and recombination. Fluorescence decays are shown for molecules having 139-bp, 146-bp, and 153-bp DNA loops. Rate constants were obtained by fitting the fluorescence decay to a system of ordinary differential equations that describe the time-dependent concentrations of reactants, intermediates and products along the intramolecular recombination pathway.^36^ The best-fit numerical solution is given by the solid curve. The fluorescence decay and fit to the data over the first 15 minutes of the recombination reaction are shown in the inset.

### Measurements of DNA helical repeat under recombination-reaction conditions

*J* as a function of the loop size *n* is a periodic function, whose amplitude, offset, and phase are not single-valued with respect to synapse geometry. In particular, interpreting the measured *J*(*n*) values in terms of a geometric model of the Cre-loxP synapse depends strongly on the DNA helical repeat, *h_0_*. Both small changes in synapse geometry and *h_0_* can lead to pronounced shifts in the theoretical dependence of *J* on loop size; indeed, accurate solutions for the geometric variables that describe the conformation of the protein-DNA synapse cannot be obtained in general without knowing *h_0_*. Additional uncertainties in *h_0_* can arise in cases where unusual buffer conditions are employed, required in this case by the need to reproduce the conditions used in previous measurements of apparent Cre-loxP association/dissociation rate constants.^37^ Possible effects of solution conditions are rarely considered or independently measured despite the established fact that counterion type and concentration measurably affect DNA helical repeat.

The present experiments were carried out under solution conditions identical to those used previously,^36, 37^ in which recombination reactions contained significant concentrations of polyethylene glycol (PEG) and glycerol (10% and 20% (w/v), respectively). Previous studies of effects of “dehydrating agents” on DNA twist showed that glycerol, as well as other non-aqueous solvents such as ethylene glycol and dimethyl sulfoxide, increase the DNA helical repeat in the range of non-aqueous solvent concentrations used in our study.^41^ We therefore carried out independent measurements of the DNA helical repeat for this series of loxP substrates under our recombination-reaction conditions.

Helical-repeat values were measured using a technique developed in our laboratory that is based on the dependence of gel-electrophoretic mobility on plasmid size for individual DNA topoisomers.^9^ The mobilities of topoisomers resolved in one-dimensional electrophoresis experiments vary discontinuously at critical values of the construct size, leading to a saw-tooth dependence on plasmid size that is highly sensitive to the helical repeat of the size-varying region. By fitting this size dependence to parameters that affect the writhe of superhelical molecules, including the torsional and bending persistence lengths of the DNA, we found that the most-probable helical-repeat value for the looped DNA segment in our recombination buffer is 10.76 (±0.05) bp turn^−1^, a significant departure from the canonical value of 10.45 bp turn^−1^.^42^ (Supplementary Figure 2).

### Fitting the experimental values of J to a structural model of the Cre-loxP synaptosome

In a previous analysis of lac repressor-operator complexes we used the helical dependence of gene repression over short curvilinear operator-operator distances to determine the most-probable geometry of the lac-repressor tetramer.^43^ The theory used to fit those data was based on a semi-analytical harmonic approximation to the free energy of DNA looping.^44^ Here we employ an extension of this approach based on normal-mode analysis (NMA) to fit a structural model of the Cre-loxP synaptosome to the experimentally measured J-factor values, *J_exp_*(*n*), where *n* is the number of base pairs in the Cre-loxP mediated DNA loop. NMA allows one to calculate the free energy and to identify the principal modes of vibration of macromolecular structures fluctuating about a mechanical-energy minimum and is mathematically equivalent to the harmonic approximation.^45, 46^

The present structural model of the Cre-loxP synaptosome regards the looped nucleoprotein complex as a Cre half-tetramer anchoring the base of a DNA loop *n* base pairs in size. DNA base pairs and individual Cre subunits are treated as a connected set of rigid bodies as described in ^44^ and ^47^. Whereas DNA base pairs and loxP-bound Cre monomers are treated as rigid entities, their respective interactions with adjacent base pairs and neighboring protein subunits are governed by purely harmonic potential energies (see Figure 3 and Methods).

**Figure 3.**
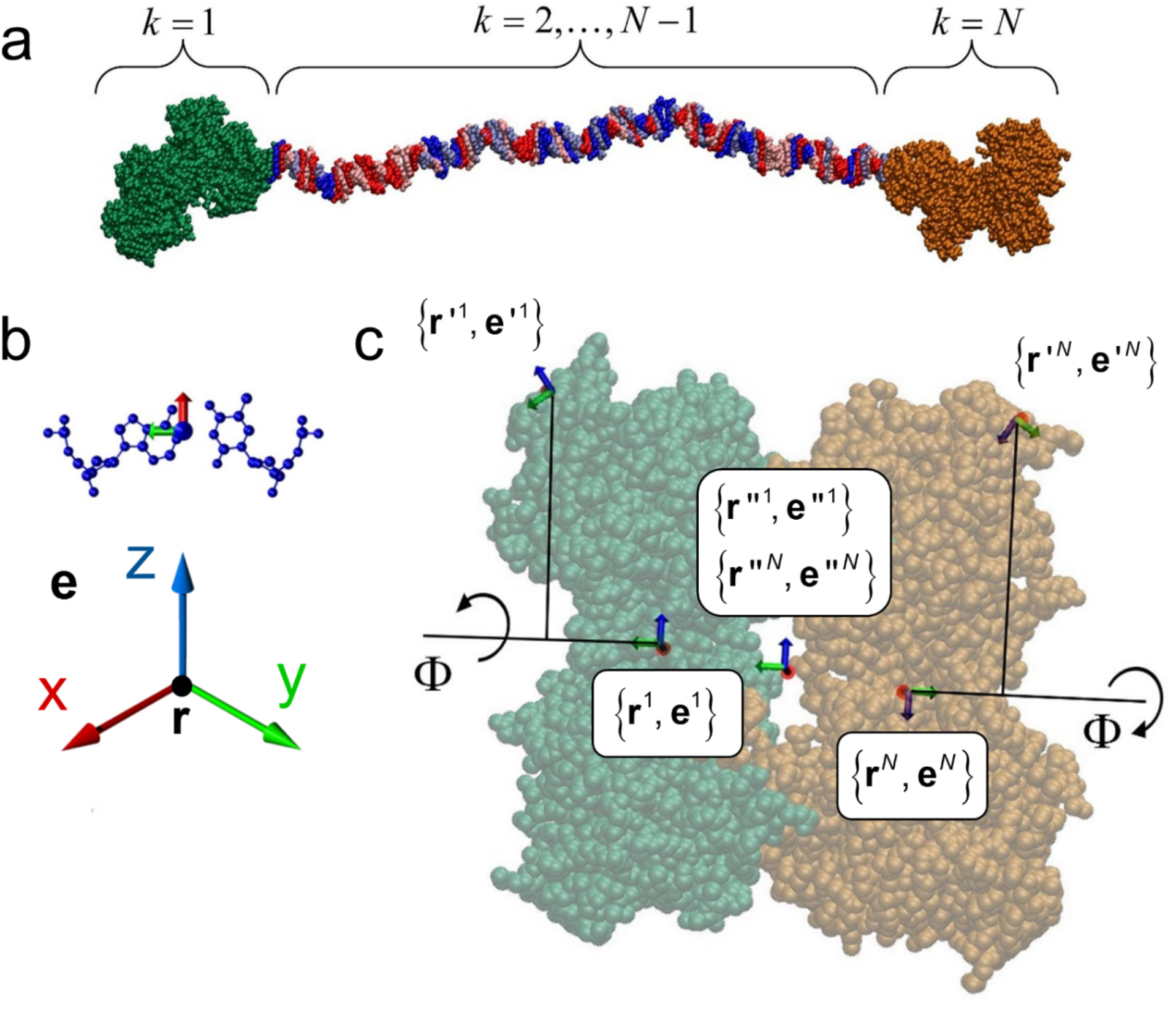
Definition of the dihedral angle between Cre half tetramers in the synapse. The geometry of the Cre-loxP synaptic complex was modified from the nearly planar crystal structure geometry by applying rotations of the Cre-bound loxP sites as shown. Frame {**r′**^1^, **e′**^1^} is rotated counterclockwise by an angle Φ about the axis 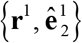 and frame {**r′**^*N*^, **e′**^*N*^} is rotated counterclockwise by the same angle Φ about the axis 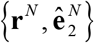, increasing the dihedral angle of the loop ends in the protein synapse by 2Φ.

In recent work we developed a general method, termed TI-NMA, for computing the conformational free energy of DNA tertiary structures and nucleoprotein assemblies based on a combination of thermodynamic integration (TI) and NMA.^45^ By augmenting NMA with TI it is possible to determine the conformational free energies of a wide range of nucleoprotein structures (and those of other macromolecules) across length scales.

Importantly, the TI-NMA method also provides a quantitative estimate of the deviations from exact results for free energies computed from NMA (or the harmonic approximation) exclusively. For DNA molecules on the length scale considered here (≈ 200 bp) the deviation of the free energy computed by NMA alone from exact values amounts to less than the thermal energy, *k_B_T* (where *k_B_* is the Boltzmann constant and *T* = 300 K is the temperature); in terms of *J* this error is approximately within a factor of two.^46^ We account here for this theoretical deviation as well as potential uncertainties in the exact value of *K_b_* by introducing an adjustable multiplicative constant *α* relating computed J-factor curves to those obtained by applying NMA to our structural model, *i.e*., *J*_comp_(*n*) = *αJ*_NMA_(*n*). ^22^ In order to fit the computed J-factor curves to the experimental J-factor values we took both the dihedral half-angle Φ subtended by the Cre monomers (Figure 3) and an associated elastic-energy constant *k_r_* (in units of *k_B_T* ·rad^−2^) as adjustable parameters.

Figure 4a shows the two-dimensional map of residuals 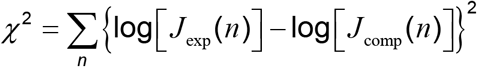 as a function of Φ and *k_r_* for wild-type Cre-mediated looping. In this calculation we optimized the parameter *α* in the relation *J*_comp_(*n*) = *αJ*_NMA_(*n*) to minimize *χ*^2^ for each pair of values (Φ, *k_r_*). The DNA properties used in the calculation were based on uniform (non-sequence-dependent) helico-elastic parameters, including the measured sequence-averaged helical repeat of 10.76 bp turn^−1^. The *χ*^2^ plot exhibits a clear minimum at Φ = 33° and *k_r_* = 2.5 *k_B_T* rad^−2^ (Figure 4a). Figure 4b shows experimental *J_exp_* values along with computed J-factor curves corresponding to the envelope of optimal Φ and *k_r_* values: +27° ≤ Φ ≤ +33° and 2 *k_B_T* rad^−2^ ≤ *k_r_* ≤ 3 *k_B_T* rad^−2^. In contrast, the helical dependence of *J* for Φ = 0° and Φ = −33° (achiral planar and left-handed geometries, respectively) shown in Figure 4b do not provide satisfactory fits to the experimental data.

**Figure 4.**
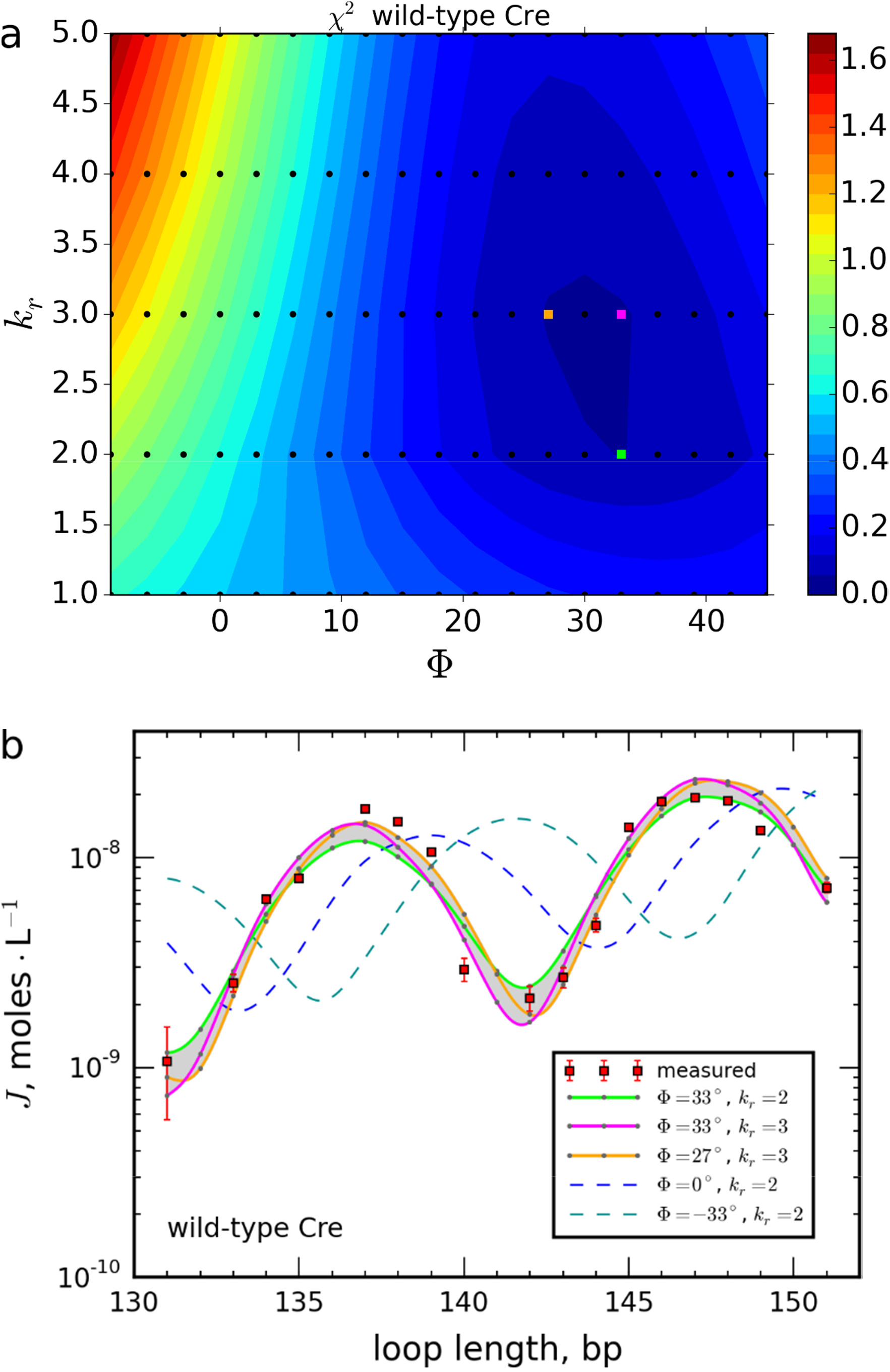
Contour map of solutions and optimal fits of calculated J-factor curves to measured J-factor values for wild-type Cre-mediated looping. (**a**) Contour plots of the mean-square error, *MSE*, for fitting of experimental J factors to NMA-computed values for the wild-type Cre synapse as a function of the dihedral angle Φ and rotational flexibility constant *k_r_* for the pair of Cre half-tetramers (see Methods). **(b)** Measured values of *J* for wild-type Cre along with an envelope of optimally fitted J-factor curves, corresponding to values (Φ, *k_r_*) indicated by the three highlighted points in the contour map shown in **(a)**. Each data point is the average of at least three independent measurements and error bars indicate ±1 standard error. The envelope of optimal Φ values spans the range 27° (orange) ≤ Φ ≤ 42° (green) with 2 ≤ *k_r_* ≤ 3 *k_B_T*·rad^−2^. A particular solution in this range for Φ = 33° with *k_r_* = 3 *k_B_T*·rad^−2^ is shown in magenta. Calculated J-factor curves for Φ = 0° (blue dashed curve) and Φ = −33° (cyan dashed curve) with *k_r_* = 2 *k_B_T*·rad^−2^, provided for comparison, do not provide satisfactory fits to the experimental data.

## Discussion

Although development of theories for DNA-loop formation (and DNA cyclization as a special case) has progressed over more than twenty-five years, application of the theories to analyze experimental data have not been widespread. One potentially significant barrier to wider use of loop-closure kinetics in solution is a scarcity of methods for monitoring the kinetics of loop formation with adequate temporal resolution to provide reliable values of rate constants. Here we employ a FRET-based measurement of recombination-site synapsis at sufficient temporal resolution to investigate the intramolecular synapsis of two loxP sites tethered by short (≲ one persistence length) DNA segments. The use of small looped DNAs in such assays allows the geometry of the looped recombinase-DNA complex to be determined from J-factor measurements with high sensitivity. This approach can be a valuable complement to the analysis of nucleoprotein assemblies by topological methods, which generally remain ambiguous with respect to geometric details.

Here we focus on a particular asymmetry in the synaptic complexes of tyrosine site-specific DNA recombinases (λ Int, Cre, and Flp). A previous topological study of tyrosine recombinases, including Cre, revealed an excess of (+)-noded knotted recombinant DNA products over (-)-noded topologies.^29^ This excess implies the existence of a productive right-handed recombination intermediate; however, no available crystallographic structure for any of these recombinases with their DNA target sites shows significant right-handed chirality. The data presented here provide the first direct evidence for a right-handed recombination intermediate in solution and also support apparent asymmetries in the arrangement of recombination sites observed by atomic-force and electron microscopy.^30, 50^ It is difficult to conceive of a mechanism whereby product chirality arises from solely non-productive conformational preferences. Therefore, we conclude that the chiral intermediate described here likely has an active role in the recombination pathway.

Our method considers the dynamic behavior of Cre synaptic complexes rather than a single static, or average, structure. An inherent characteristic of ensemble-based kinetic methods is that they typically probe only the intermediate species with the longest lifetime. Thus, it is possible that other synapse geometries with shorter lifetimes, including achiral crystallographic structures, may exist along the recombination pathway. A single-molecule FRET analysis based on various crystallographic synaptic complexes identified a subset of possible intermediate conformations that may characterize distinct steps in the recombination pathway^51^. Due to the inherent difficulties of obtaining absolute distance measurements from FRET-efficiency values.^36, 52, 53^ it is difficult to assign a particular geometry to the observed intermediates and the possibility of one or more productive non-planar intermediate structures should not be ruled out.

Finally, to resolve potential roles of specific mechanistic steps in the Cre-recombination pathway, we analyzed the loop-closure kinetics for a subset of loxP substrates using a mutant form of Cre (R101A), which is defective in isomerization and resolution of the central Holliday-junction intermediate. Our analysis shows that the mutant synapse is also right-handed with a somewhat larger dihedral angle (Φ ≈ 45°) relative to the wild-type complex (Fig. 5). In the wild-type reaction we speculate that, if there is sufficiently rapid averaging of a structure with a larger Φ value at the isomerization step together with a structure that is less chirally biased (*e.g.*, Φ ≈ 0), this could explain the overall smaller value of Φ (33°) measured for the wild-type recombinase.

**Figure 5.**
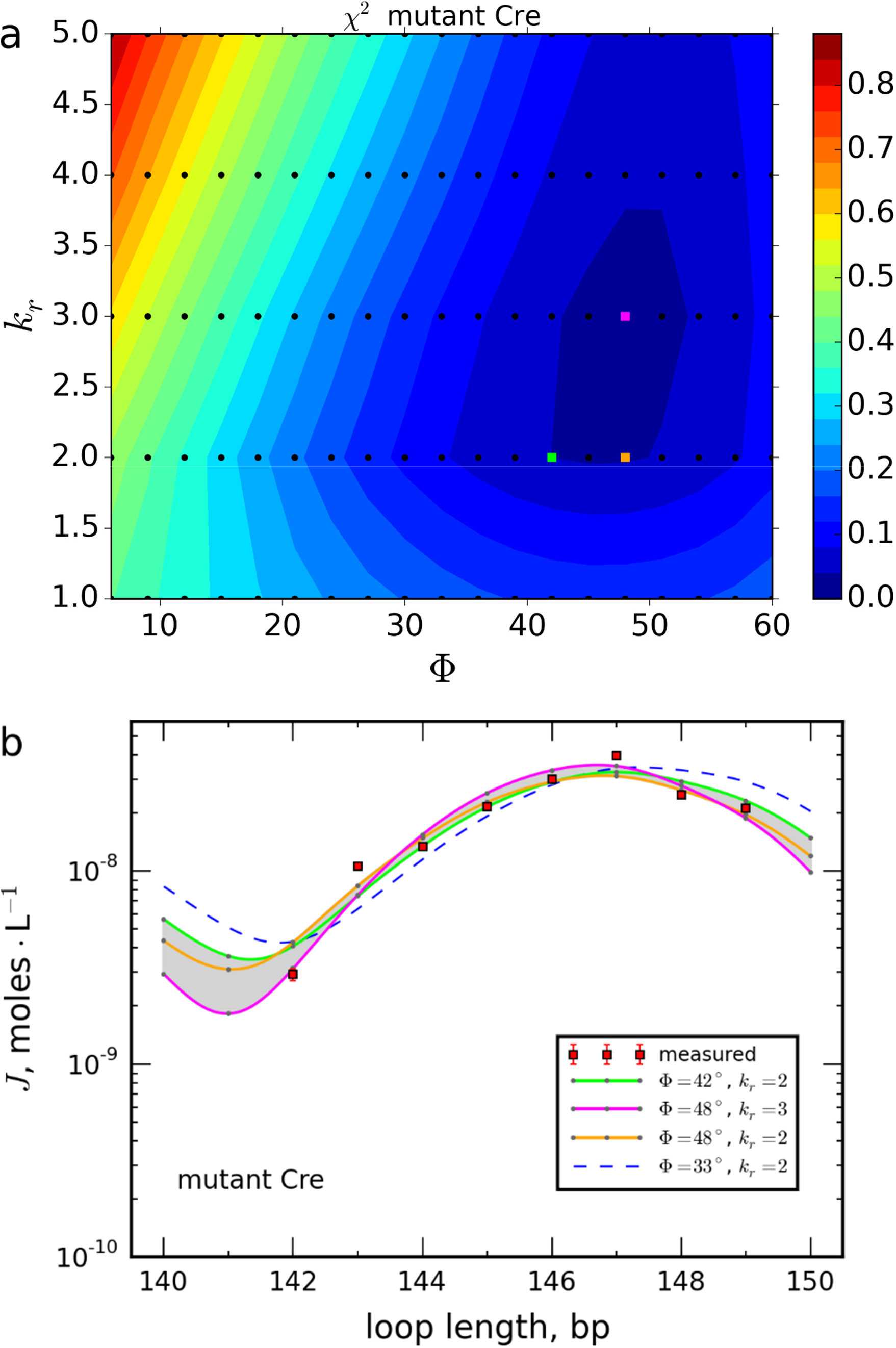
Contour map of solutions and optimal fits of calculated J-factor curves to measured J-factor values for Cre-mediated looping by the R101A mutant Cre protein. The R101A mutation strongly reduces the protein’s Holliday-junction resolution activity, promoting the accumulation of junction-containing synaptic complexes. Contour plots **(a)** and optimally fitted J-factor curves **(b)** for mutant Cre were obtained as described in Figure 4. Optimal fits were obtained for 42° (green) ≤ Φ ≤ 48° (orange) and *k_r_* = 2 *k_B_T*·rad^−2^. A particular solution for Φ = 48° with *k_r_* = 3 *k_B_T*·rad^−2^ is also shown in magenta. The J-factor dependence for Φ = 33° and *k_r_* = 2 *k_B_T*·rad^−2^, which lies within the best-fit envelope of the wild-type protein, is shown for comparison (blue dashed curve).

The existence of a stable conformational intermediate with significant right-handed chirality leads to symmetry breaking in the distribution of recombination-product topologies. It may also reflect biological asymmetries that arise from the right-handed geometry of the DNA double helix. Packing of DNA helices at high densities such as in some crystal structures,^54–56^ have biases for right-handed configurations that are sterically favorable due to reciprocal fitting of the DNA backbone and duplex major groove. Right-handed duplex geometry also contributes to the chiral organization of DNA in viral capsids^57^ and some liquid-crystalline phases.^58, 59^ Finally, crystal structures of isolated HJs (in the absence of junction-binding proteins) seem to invariably adopt right-handed “stacked-X” structures with interduplex angles comparable to the values measured in this work.^60, 61^ The consensus view is that HJs are highly dynamic entities, a conclusion that is supported by the kinetic analysis presented here.

## Methods

### Plasmid DNAs

A series of plasmids pCS2DloxP(*n*) were constructed by inserting DNA fragments with lengths (*n* − 34) bp, which were derived by PCR amplification from the bacteriophage lambda genome, between the *Pst*I and *Not*I sites of pCS2DloxP.^36^ For example, plasmid pCS2DloxP(130) was constructed by inserting a 96-bp DNA fragment between the two loxP sites as described. Cloning steps were confirmed by didexoynucleotide sequencing. Plasmids were propagated in *E. coli* HB101 cells and isolated in quantity by using a Promega Wizard Plus Megaprep purification kit (Promega Corp., Madison, WI).

### Fluorescently labeled DNAs

Donor and acceptor fluorophores were incorporated into DNA fragments using PCR. Plasmids from the pCS2DloxP(*n*) family were linearized by ScaI treatment and used as templates. Atto 647N- and Atto 594-labeled oligonucleotides were used as forward and reverse primers, respectively. The primers were purchased from IBA (Göttingen, Germany) and were purified twice by reverse-phase HPLC. PCR reactions were carried out using *Taq* DNA polymerase and dNTPs from New England Biolabs (Ipswich, MA) (see Supplementary Figure 1). Excess primers were removed from the PCR products using NucleoSpin silica-membrane columns (Clontech, Mountain View, CA) and the products were further purified in 3% agarose-TBE gels and subsequently reisolated using gel-purification kits (Clontech).

### Protein Purification

The expression vector for the His-tagged wild-type Cre_WT_ (pET28b-His6Cre) was a gift from Dr. Enoch Baldwin at UC-Davis.^22, 26^ The vector for the mutant Cre_R101A_ protein (pLC101A) was a gift from Dr. Paul Sadowski at the University of Toronto.^62^ Both proteins were expressed from BL21(DE3) cells bearing the corresponding plasmids and were purified according to.^63^ Partially purified lysates were loaded onto equilibrated cobalt His-TALON columns (Clontech, Mountain View, CA). The column was washed with 40 to 50 volumes of wash buffer (50 mM NaH_2_PO_4_, 300 mM NaCl, 10 mM imidazole; pH 7.8). The Cre protein was eluted using an imidazole gradient (10 mM - 150 mM). All of the eluted fractions were analyzed by SDS-PAGE electrophoresis with Coomassie and AgNO_3_ staining. The protein preparations were estimated to be over 90% homogenous. Cre-containing fractions were pooled and dialyzed against (20 mM Tris-Cl, 700 mM NaCl, 0.5 mM EDTA, 2 mM DTT and 0.05% (w/v) sodium azide; pH 7.8). Protein was concentrated using a centrifugal concentrator with an MWCO below 15 kDa (Millipore). The concentrated protein was sub-aliquoted, flash frozen, and stored at −80 °C.

### FRET-based recombination-kinetics assays and analysis

Kinetics measurements were carried out using the pure fractions of Cre_WT_ and CreR_101A_ under conditions identical to the intramolecular-recombination assays described in ^36^. We employed numerical methods to extract rate constants for fundamental steps in the recombination pathway corresponding to synapse formation and dissociation. The modeled recombination pathways for both intermolecular and intramolecular recombination mechanisms and additional details of the curve-fitting procedure are also given in ^36^. Curve-fitting routines were implemented in Matlab and used the functions *lsqcurvefit* for non-linear least-squares fitting and *ode15s* to solve the initial-value problem for the systems of ordinary differential equations. Analysis programs are available upon request.

### Computational modeling of the looped synaptic complex

A Cre-bound DNA synapse was modeled as a chain of *N* rigid bodies labeled *k* = 1,…,*N*. The terminal rigid bodies *k* = 1, *N* represent Cre-bound loxP sites and the remaining rigid bodies *k* = 2,…, *N* − 1 represent single DNA base pairs spanning the Cre-bound loxP sites (Supplementary Figure 3). In terms of loop size, *n*, which is defined as the curvilinear distance in base pairs between centers of the flanking loxP sites, *N* = *n* − 32. Embedded within each rigid body is a body-fixed reference frame with positions and orientations {**r**^*k*^, **e**^*k*^}. The boundary conditions for base pairs *k* = 2, *N* − 1 adjacent to the Cre-bound loxP sites are described by additional body-fixed auxiliary frames {**r′**^1^, **e′**^1^}, {**r′**^*N*^, **e′**^*N*^}, associated with the central DNA base pairs embedded within the Cre-bound loxP sites. The interaction between loxP sites in a Cre-mediated DNA loop is further described in terms of body-fixed auxiliary frames {**r″**^1^, **e″**^1^}, {**r″**^*N*^, **e″**^*N*^} embedded in the loxP sites. The equilibrium shape (geometry) of the individual protein-DNA complexes and of the overall synaptic complex is fully characterized by the displacements of the auxiliary frames {**r′**^*k*^, **e′**^*k*^}, {**r″**^*k*^, **e″**^*k*^} relative to the reference frames {**r**^*k*^, **e**^*k*^} for *k* = 1, *N*.

The potential energy of a specific configuration of the Cre-bound DNA complex is given by

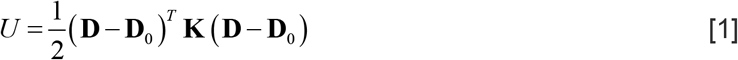

where the vector **D** contains the 6*N* displacements between neighboring rigid bodies, **D**_0_ are the mechanical-equilibrium displacements for the linear form of the complex, and **K** is a 6*N* × 6*N* stiffness matrix. Assuming only nearest-neighbor interactions, the displacement between any pair of frames *k* and *k* + 1 is parameterized in terms of 6 local displacement coordinates (shift, slide, rise, tilt, roll, twist), which were calculated as detailed in ^47^. We took the helical parameters for **D**_0_ to be those of canonical B-form DNA with the exception of the twist (*i.e*., shift = slide = 0, rise = 0.34 nm, tilt = roll = 0, twist = 2*π*/*h*_0_ = 33.46° with *h*_0_ = 10.76 bp turn^−1^). A sequence-averaged stiffness matrix was obtained as 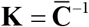 where 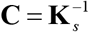 is the covariance matrix of the stiffness matrix **K**_*s*_ obtained in ^64^ and the overbar denotes an average over the 10 independent base steps. Auxiliary frames {**r″**^*N*^, **e″**^*N*^}, {**r″**^1^, **e“**^1^} were defined such that at mechanical equilibrium {**r″**^*N*^, **e″**^*N*^} = {**r″**^1^, **e″**^1^}, i.e., 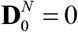. The stretching and rotational flexibility between the Cre-bound loxP sites was described by force constants *k_s_* and *k_r_*, respectively. We used *k_s_* = 5 *k_B_T*Å^−2^ and *k_r_* was determined from fitting the model to experimental values of the J factor.

Initial positions and orientations of rigid bodies *k* = 1, *N* describing Cre-bound loxP sites were obtained from the Protein Data Bank (PDB) for the Cre-loxP synaptic complex (PDB ID 5CRX). Coordinates of auxiliary frames {**r′**^1^, **e′**^1^}, {**r′**^*N*^, **e′**^*N*^} were deduced using the software package 3DNA,^65^ and coordinates of auxiliary frames {**r″**^1^, **e″**^1^}, {**r″**^*N*^, **e″**^*N*^} were obtained from the crystal structure of the complete synaptic complex (2 loxP + Cre tetramer). In our study, the geometry of the Cre-bound loxP sites was allowed to vary via a single rotational degree of freedom (Supplementary Figure 3). Frame {**r′**^1^, **e′**^1^} was permitted to rotate by an angle ±Φ about the axis 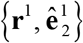 and frame {**r′**^*N*^, **e′**^*N*^} was rotated by the same angle ±Φ about the axis 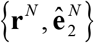. These rotations therefore increase the dihedral angle of the Cre-mediated loop at the protein synapse by 2Φ. By convention, a positive rotation of angle 2Φ corresponds to a righthanded crossing of the loxP sites.

Minimum-energy configurations of a Cre-bound DNA complex were obtained by using unconstrained minimization of the potential energy *U* in Eq. (1) with a trust-region and normal-mode analysis (NMA) was carried out as described in ^45^ and ^46^. Theoretical J factors were calculated as the ratio *J* = *K_c_*/*K_b_* of intramolecular and bimolecular equilibrium constants using a generalization of the NMA method in ^46^. In particular, *K_b_* was calculated by applying NMA to a pair of bound and unbound rigid bodies describing Cre-bound loxP sites in the absence of an intervening DNA loop. J values were calculated for each minimum-energy configuration having preferred (lowest energy) linking number *Lk*_0_. J factors for topoisomers with *Lk* = *Lk*_0_ ± 1,2,… were approximated by fitting calculated J-factors with preferred linking number *Lk*_0_ to the function *f*(*x*) = exp[−*c*_2_(*x* − *c*_1_)^2^ + *c*_2_] and extrapolating beyond *Lk*_0_. Final J-factor values, accounting for multiple topoisomers, were calculated by summing *J* for all linking numbers. For illustration, J-factors calculated for a Cre-mediated DNA loop are shown in Supplementary Figure 4 for a range of DNA loop lengths 120-164 base pairs, using *k_s_* = 5 *k_B_T*Å^−2^, *k_r_* = 5 *k_B_T*·rad^−2^, and DNA parameters **D**_0_, **K** described above. The equilibrium protein geometry was generated from the crystallographic structure of the Cre-loxP synaptic complex (PDB: 5CRX).^66^

## Acknowledgments

Authors would like to thank Heidi Kim for her assistance in plasmid purification. This manuscript is dedicated to the memories of Nick Cozzarelli and Don Crothers.

## Author contributions

SDL conceived of the study, supervised the overall project, and wrote the manuscript. MS performed all experiments and wrote the manuscript. MS and SDL carried out J-factor computations and analyzed data. SMG programmed and ran J-factor computations with guidance from AH, who also wrote portions of the manuscript. AV provided preliminary data and aided MS at early stages of project development. RZ participated in data analysis and assisted with editing of the manuscript.

